# A dynamic redox switch turns TRC40 into a chaperone protecting human cells against ATP-depleting, oxidative stress

**DOI:** 10.1101/2024.07.10.602939

**Authors:** Bianca Dempsey, Risai Dubrall, Olivia Chan, Kim Jasmin Lapacz, Jan Riemer, Ursula Jakob, Kathrin Ulrich

## Abstract

Oxidative stress represents a major challenge for cellular proteostasis. The accumulation of reactive oxygen species, such as hydrogen peroxide, impairs the fidelity of protein biosynthesis and causes non-specific oxidative protein modifications and aggregation. This situation is further aggravated by the oxidative stress-mediated drop in cellular ATP levels, which reduces the activity of ATP-dependent chaperones and proteases. We now demonstrate that to cope with oxidative unfolding stress, human cells rely on the moonlighting function of TRC40, which turns from an ATP-dependent targeting factor into an ATP-independent chaperone upon oxidation. Controlled by a highly conserved redox switch, oxidized TRC40 forms chaperone-active tetramers and high-molecular complexes which prevent the aggregation of unfolding proteins. Acute oxidative stress leads to the reversible formation of distinct TRC40 foci, associated with the canonical chaperones Hsp70 and Hsp110, suggesting a role of these stress-induced structures in recovering and sorting aberrant proteins. Consistently, we discovered that TRC40 is essential upon ATP-depleting, oxidative stress conditions to counteract the accumulation of mis- and unfolded proteins, which are rapidly cleared in an TRC40-dependent manner. Our data reveal that TRC40 is an integral part of protein quality control and support its role as a triaging factor in cellular proteostasis.

## INTRODUCTION

Maintaining a healthy proteome is essential to accomplish all aspects of biological functions and ensure cellular integrity. From the biosynthesis of a polypeptide chain at the ribosome to the coordinated degradation of mature proteins, a complex and highly dynamic network of proteostasis factors manage and control protein folding, targeting, complex assembly and disassembly as well as clearance of aberrant proteins^1–4^. Oxidative stress represents a major challenge for these systems. Independent of the source, an increase in reactive oxygen will cause unspecific protein oxidation endangering folding and function of proteins^5,6^. In addition, oxidative stress results in a drop of cellular ATP levels, which impairs ATP-dependent chaperones and proteases and thus, increases the risk of potentially toxic protein aggregation^7–9^. The oxidative activation of ATP-independent chaperones serves as a first line of defense to prevent wide-spread protein aggregation^10–12^. Since the discovery of Hsp33 as the first redox-regulated chaperone in *E. coli* ^13^, stress-activated chaperones have been shown to be essential for mediating resistance towards environmental challenges in various organisms^14–18^.

The cytosolic protein Get3 has been identified to serve analogous to Hsp33 as a redox-regulated chaperone in yeast. Get3 is a dual function protein which under non-stress conditions, acts as ATPase in the <u>G</u>uided <u>E</u>ntry of <u>T</u>ail-anchored proteins (GET) pathway, facilitating the post-translational targeting of tail-anchored membrane proteins to the ER^19,20^. Upon oxidation, Get3 forms chaperone-active oligomers that effectively prevent protein aggregation^21–23^. The chaperone function has been shown to be essential and sufficient to rescue the oxidative stress-induced phenotypes of a *get3*-deletion strain^22^. We recently demonstrated that the functional switch into an ATP-independent chaperone is induced upon nucleotide release followed by the oxidation of a highly conserved CXC motif^23^. Upon restoring reducing conditions and ATP level, Get3 transfers bound proteins to the ATP-dependent Hsp70/Hsp40 system for refolding^23^. In cells, chaperone-active Get3 forms defined foci, colocalizing with Hsp70/Hsp40 as well as other ATP-dependent chaperones^21,22^. Get3 mutant variants lacking the thiol switch fail to release bound clients and do no longer dissociate from foci. Hence, the functional switch is required for both, the stress-specific activation of the chaperone function and the coordinated release of client proteins once non-stress conditions have been restored^23^.

Although Get3 is highly conserved in eukaryotes and its ability to interact with unfolding proteins appears to be an ancient feature^24^, it is still unknown whether the redox-regulated chaperone function is conserved in mammalian cells and the extent to which it protects cells against oxidative unfolding stress. TRC40 (also known ASNA1) is the metazoan homolog of Get3 (Figure S1A and B) and has been well-characterized as part of the mammalian transmembrane recognition complex (TRC) pathway, targeting tail-anchored membrane proteins to the ER^25–27^. The deletion of TRC40 is embryonically lethal^28^ and conditional knockout approaches suggest a cell type-specific function of TRC40 in tissue development^29^. Numerous structural and functional studies contributed to the current model of the TRC pathway, which represents a robust route for tail-anchored proteins to the ER^20,27,30,31^. Surprisingly, however, only a subset of tail-anchored proteins relies on the TRC pathway while most can utilize alternative routes^29,32^. These findings raise the question whether TRC40 serves an additional cellular function as a molecular chaperone – an activity that would become especially important upon oxidative stress when ATP-dependent protein insertion is impaired and unfolding intermediates accumulate in the cytosol.

Here, we demonstrate that TRC40 functions as a redox-regulated chaperone, which is exceptionally sensitive towards oxidative activation once it is in its nucleotide-free state and efficiently prevents aggregation of unfolding protein intermediates. In human cells exposed to ATP-depleting, oxidative stress, TRC40 forms reversible foci which spatially overlap or associate with the ATP-dependent chaperones Hsp70 and Hsp110. Presence of TRC40 is crucial to cope with H_2_O_2_-induced protein aggregation, revealing its role as a redox-regulated factor of the proteostasis network preventing protein aggregation upon oxidative stress.

## RESULTS

### TRC40 moonlights as a redox-regulated chaperone

Multiple structural and functional studies revealed that TRC40 mediates the ATP-dependent binding and targeting of tail-anchored proteins in mammalian cells^25–27^. Yet, all *in vitro* studies were conducted under reducing conditions, where the protein is present as a non-covalently linked dimer with moderate ATPase activity^30,33–36^. To investigate whether TRC40 is sensitive towards oxidation and potentially switches into a molecular chaperone, we used zebrafish TRC40, which shares 93.8% identify with human TRC40 (Figure S1B), but can be stably expressed and purified from *E. coli* ^37^ (Figure S1C). We tested the chaperone activity of TRC40 by monitoring its influence on the aggregation of chemically denatured citrate synthase (CS). Indeed, treatment with 2 mM H_2_O_2_ and CuCl_2_ for 10 min at 37°C activated the chaperone function of TRC40, as indicated by the strongly reduced CS aggregation in the presence of a 10-fold molar excess of the TRC40 (Figure 1A and B). We set this observed chaperone activity to 100% and applied the same conditions in all following chaperone assays. As previously observed with yeast Get3^22,23^, we found that the oxidative activation of TRC40’s chaperone function reversibly inactivated its ATPase activity (Figure 1C). Incubation of oxidized TRC40 with DTT in the presence of Mg-ATP inactivated the chaperone function and largely restored ATPase activity (Figure 1B and C). Non-reducing SDS-PAGE analysis revealed that TRC40 oxidation results in various species with intra- and intermolecular disulfide bonds, which were fully reversed upon adding reducing agents (Figure 1D). Size exclusion chromatography demonstrated that reduced TRC40 is present as a dimer and forms tetramers and higher oligomeric species upon oxidation, like yeast Get3 with tetramers being the smallest chaperone-active unit (Figure 1E)^22,23^. In addition, we observed a peak eluting slightly later than the reduced TRC40 which most likely represents a species with intramolecular disulfide bonds inducing a more compact conformation. This species is also visible as a faster migrating band in the non-reducing SDS gel (Figure 1D).

**Figure 1:**
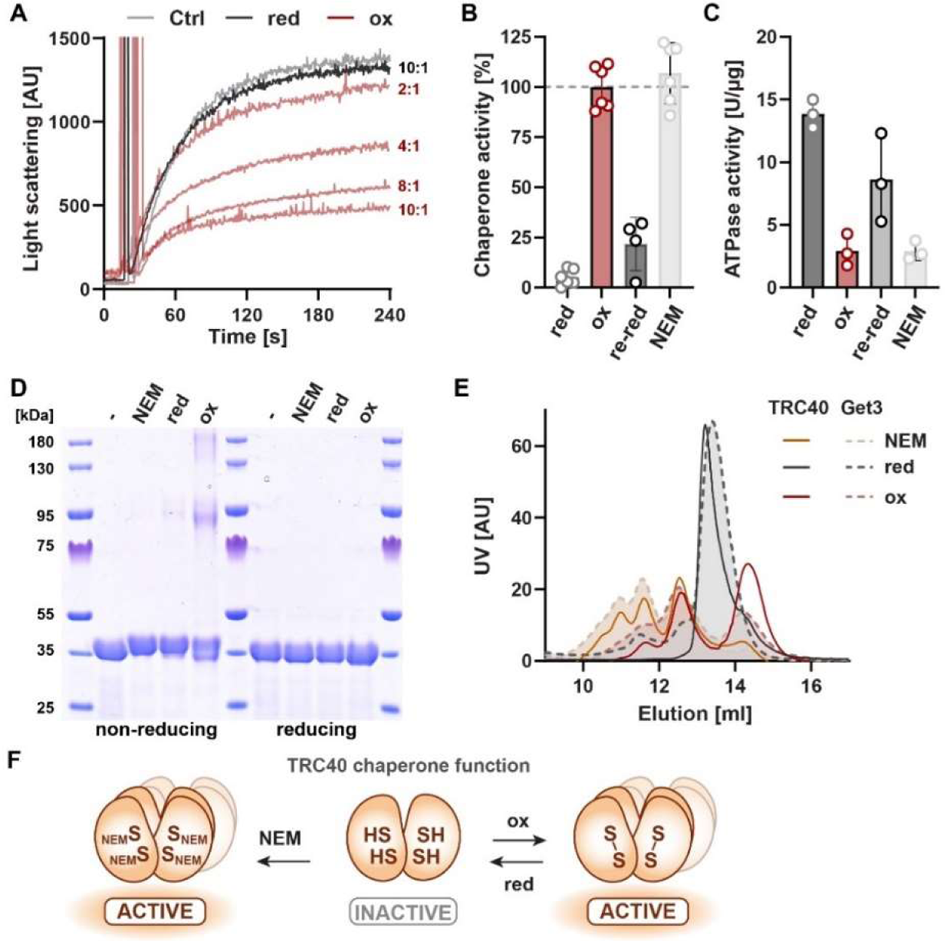
Redox-dependent functional switch of recombinant TRC40. Reduced TRC40 (**red**) was treated with 2 mM H_2_O_2_/50 µM CuCl_2_ for 10 min at 37°C to generate oxidized TRC40 (**ox**). To re-reduce TRC40 (**re-red**), oxidized TRC40 was incubated with 5 mM DTT and 2 mM Mg-ATP for 4 h at 30°C. For thiol blocking, reduced TRC40 was incubated with 10 mM with *N*-ethylmaleimide (**NEM**) for 30 min at RT. (**A**) Aggregation of chemically denatured citrate synthase (CS) at 30°C in the absence (**Ctrl**) and presence of reduced or oxidized TRC40 in molar excess as indicated. (**B**) Relative chaperone activity of TRC40. Light scattering of CS alone (Ctrl) was set to 0% and the signal observed in the presence of a ten-fold molar excess of oxidized TRC40 was set to 100%. (**C**) Specific ATPase activity of TRC40. (**B**) and (**C**) show the mean ± standard deviation (SD) of at least three independent experiments. (**D**) SDS-PAGE and Coomassie staining of TRC40 under non-reducing and reducing conditions. (**E**) Size exclusion chromatography of TRC40 and Get3 upon indicated treatments. (**F**) Scheme of the conformational change into the chaperone-active form induced by reversible thiol oxidation or irreversible alkylation. The expression and purification of recombinant TRC40 is shown in Figure S1.

One peculiarity of Get3’s thiol switching mechanism is that not the formation of a disulfide bond but the removal of the thiol groups of the conserved CXC motif is responsible for the formation of chaperone-active oligomers^23^. To test whether this is also the case for TRC40, we treated reduced TRC40 with *N*-ethylmaleimide (NEM), a thiol-reactive agent that irreversibly alkylates cysteines residues. Indeed, we found that thiol blocking was sufficient to induce the functional switch of TRC40 into its chaperone-active and ATPase-inactive form (Figure 1B and C). Although NEM prevented the formation of disulfide bonds (Figure 1D), the elution profile of NEM-treated TRC40 displayed the same pattern as oxidized TRC40 (Figure 1E). These results showed that the loss of thiols, either upon reversible oxidation or irreversible thiol blocking, induces the functional and structural switch (Figure 1F). Hence, TRC40 oligomerization does not depend on intermolecular disulfide bonds but is mediated via non-covalent interactions of the oxidized protein.

### TRC40 is exquisitely sensitive towards oxidative activation

Our previous work on Get3 revealed that the oxidation of the redox-sensitive CXC motif is strongly accelerated when Get3 is in its nucleotide-free state^23^. To investigate the role of nucleotide binding in TRC40 activation, we treated the recombinant protein with H_2_O_2_ in the absence of CuCl_2_, which substantially slows down thiol oxidation^22^. Under these conditions, we found only partial activation of the chaperone function, likely due to the remaining MgATP, which effectively prevents H_2_O_2_-mediated chaperone activation (Figure 2A). Consistently, the addition of excess Mg-ATP completely prevented the oxidative activation of the chaperone, whereas the pre-treatment of TRC40 with apyrase, an enzyme catalyzing the degradation of ATP and ADP to AMP, fully activated the chaperone function upon adding H_2_O_2_.

**Figure 2:**
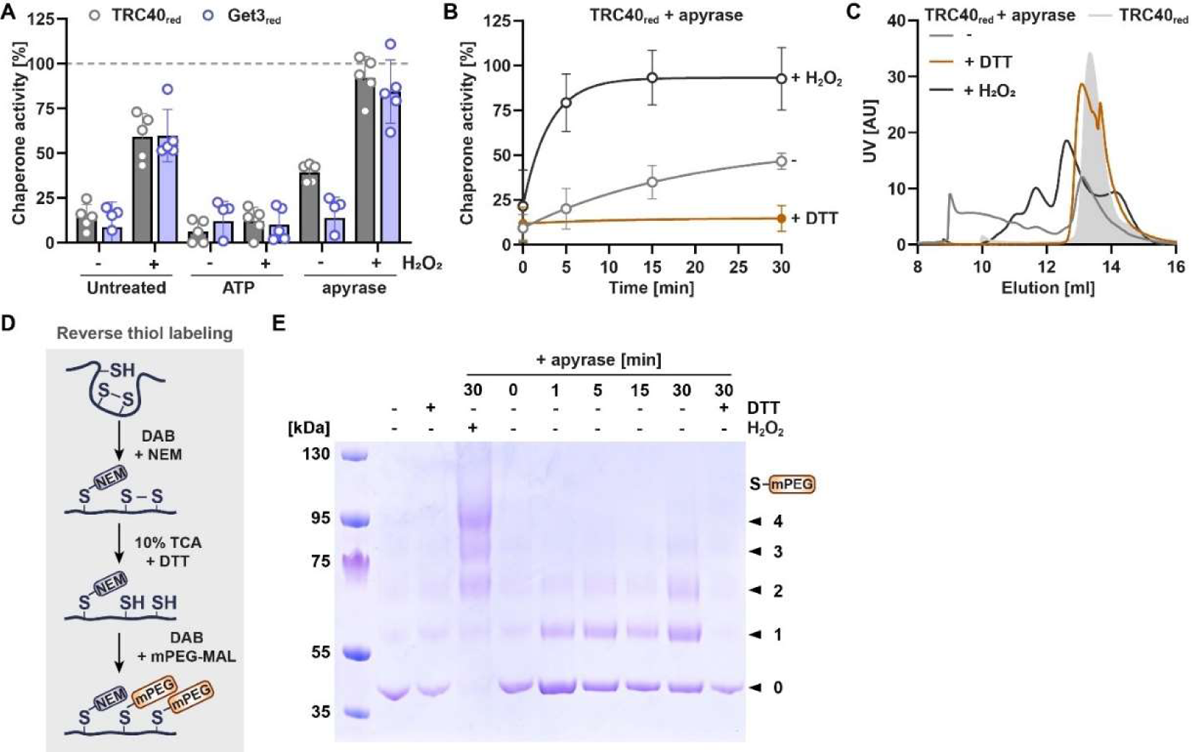
Oxidative chaperone activation of TRC40 depends on its nucleotide-binding state. (**A**) Reduced TRC40 and Get3 (untreated) were pre-incubated with either 2 mM Mg-ATP or 5 U apyrase for 15 min at 30°C and then treated without (-) and with (+) 5 mM H_2_O_2_ for 15 min. Chaperone activity was measured as described in Figure 1. (**B**) Chaperone activation of apyrase-pretreated TRC40 in the absence or presence of 5 mM H_2_O_2_ or DTT. For (**A**) and (**B**), the mean ± SD of at least three independent experiments is shown. (**C**) Size exclusion chromatography of TRC40 pretreated with apyrase and incubated without or with 5 mM H_2_O_2_ or DTT for 15 min. Reduced TRC40 which was not treated with apyrase was run as a control. Structural changes depending on the nucleotide-binding state are shown in Figure S2. (**D**) Reverse thiol labeling of oxidized cysteines. Reduced cysteines were blocked with NEM in denaturing buffer (DAB). After removing excess NEM by TCA precipitation, oxidized cysteines were reduced by DTT and newly generated thiols labeled with methoxypolyethylene glycol maleimide (mPEG-MAL), causing a mass shift per modified cysteine. (**E**) SDS-PAGE and Coomassie staining of mPEG-MAL labeled TRC40.

Remarkably, and in clear contrast to Get3, apyrase treatment of TRC40 was sufficient to activate its chaperone function even in the absence of H_2_O_2_ (Figure 2A and B). Moreover, we found that apyrase treatment also shifted TRC40 from its dimeric state into higher oligomeric structures (Figure 2C). Both processes were effectively prevented by adding excess DTT, suggesting that the activation of TRC40’s chaperone function in the absence of ATP was mediated by oxidation. To visualize the oxidation state of apyrase-treated TRC40 during air incubation, we labeled reversibly oxidized cysteines with a polyethylene glycol-linked maleimide leading to a mass increase per modified cysteine residue (Figure 2D). While reduced TRC40 ran as unmodified band at about 40 kDa, H_2_O_2_ treatment caused the formation of different TRC40 species with two to four oxidized cysteines (Figure 2E). Apyrase-treated TRC40 slowly became oxidized over time, which was prevented in the presence of DTT. These results revealed that once in its nucleotide-free state, TRC40 is highly sensitive towards oxidation, which induces the chaperone-active form.

### TRC40 forms reversible foci in human cells exposed to H_2_O_2_

During proteotoxic stress conditions, unfolding protein intermediates accumulate in distinct cellular quality control compartments enriched for molecular chaperones^38^. To investigate whether TRC40 forms part of such foci, we treated HeLa cells at 60-70% confluency with 0.5 mM H_2_O_2_ for 15 min, and monitored the cellular localization of TRC40 by immunofluorescence staining. As reported previously, TRC40 displays a broad cytosolic distribution under normal conditions^32,39^, yet readily forms of distinct TRC40 foci in about 25% of cells upon H_2_O_2_ treatment (Figure 3 A and B). The foci rapidly redissolved upon washing the cells and adding fresh media. We obtained similar results in human fibroblasts (BJ cells) and human umbilical vein endothelial cells (HUVEC), indicating that the effects of oxidative stress on TRC40 localization is observed in different human cell lines (Figure S3).

**Figure 3:**
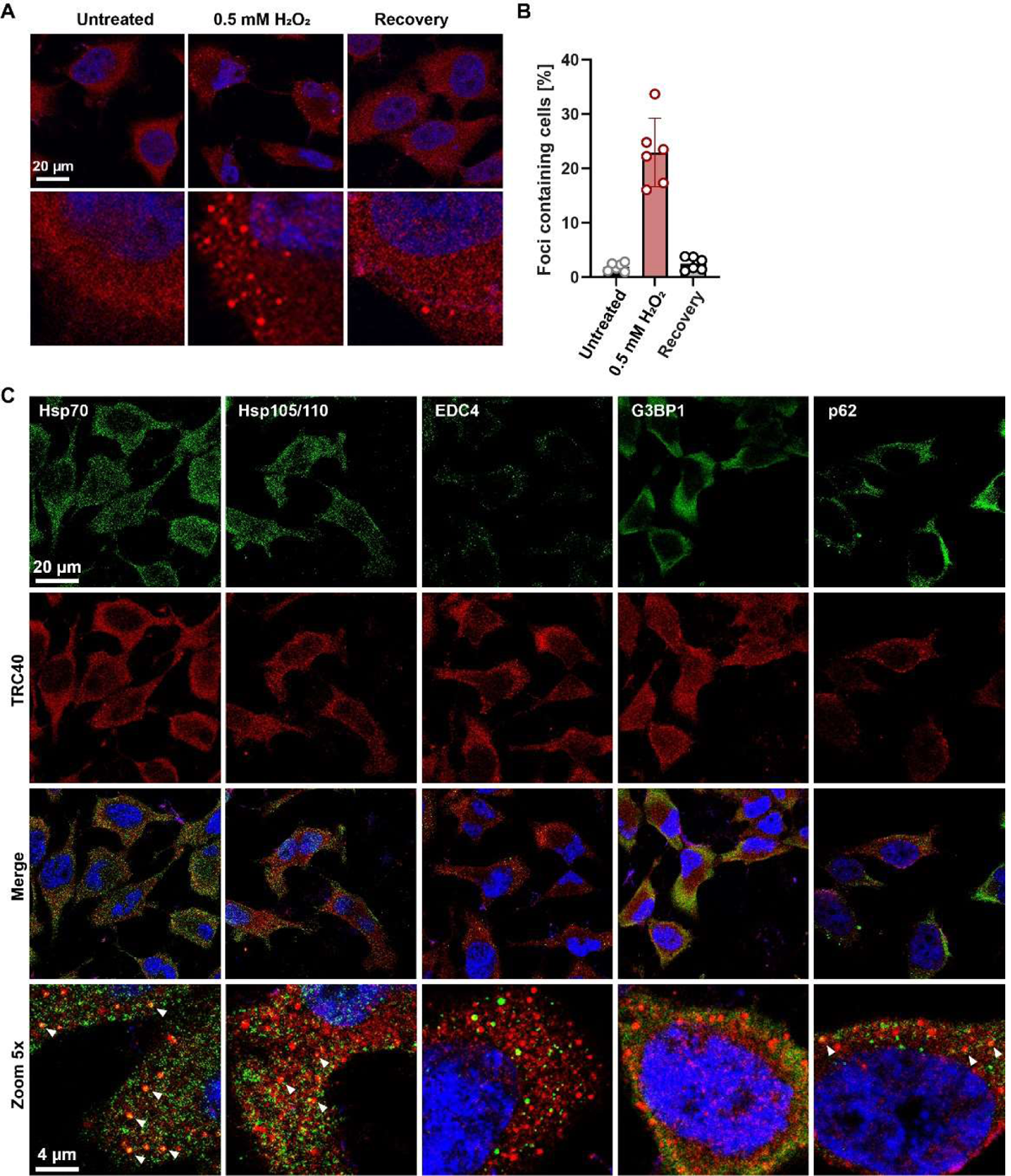
TRC40 forms dynamic punctate structures upon H_2_O_2_ exposure in HeLa cells. HeLa cells were treated with 0.5 mM H_2_O_2_ in HBSS buffer for 15 min. For recovery, cells were washed twice and supplemented with fresh DMEM. The experiment was also performed with human fibroblasts and human umbilical vein endothelial cells shown in Figure S3. (**A**) Immunofluorescence staining of TRC40. (**B**) Foci-containing cells were counted as described in Material and Methods (n = 100 cells per condition). The mean ± SD of six independent experiments is shown. (**C**) Co-immunofluorescence staining of TRC40 (in red) and the indicated proteins of interest (in green) after 15 min H_2_O_2_ treatment. Z-stack confocal microscopy images were acquired and a single focal plane of the z-stack is shown. White arrows highlight overlapping signals.

To further investigate the nature of these H_2_O_2_-induced TRC40 foci, we co-stained cells for proteins known to localize to punctate structures upon stress induction. Misfolded and stress-damaged proteins are generally shuttled to dynamic inclusions, like Q-bodies and juxtanuclear quality control compartments, which contain different molecular chaperones^38^. Indeed, we observed a clear signal for the presence of Hsp70 in TRC40-positive foci (Figure 3C), whereas co-staining with Hsp105/Hsp110 suggested a close association with TRC40 foci upon H_2_O_2_ stress. We excluded that TRC40 foci associate with RNA quality control compartments by co-staining for enhancer of decapping protein 4 (EDC4), a core P-body component^40^ or the stress granule marker G3BP1^41^ (Figure 3C). The latter was consistent with the fact that our H_2_O_2_ conditions also did not trigger the formation of stress granules in HeLa cells. Finally, we tested for co-localization with the protein sequestosome 1 (p62/SQSTM1), a selective autophagy receptor, which forms dynamic punctate structures that serve as platform for autophagosome formation in anti-oxidative stress response^42^. We found that TRC40 foci are in close proximity to the small punctate structures of p62, suggesting a potential association of TRC40 with autophagosomes.

### TRC40 is crucial to cope with H_2_O_2_-induced protein aggregation

To test whether TRC40 plays a role in protecting cells against H_2_O_2_-induced protein aggregation, we transiently transfected HeLa cells either with a scrambled, non-targeting siRNA (siCtrl) or with a siRNA targeting TRC40 mRNA (siTRC40)^32^. TRC40 protein levels were strongly downregulated 48 h post-transfection (Figure 4A and B), resulting in reduced proliferation even in the absence of oxidative stress (Figure 4C). While cell viability was only marginally affected in TRC40-depleted cells under normal conditions, these cells were more sensitive toward acute H_2_O_2_ stress compared to the control (Figure 4D). Using a genetically encoded, fluorescent ATP sensor^43^, we found that cellular ATP level rapidly dropped after treating cells with 0.5 mM H_2_O_2_. Based on TRC40’s ATP-independent chaperone function, which becomes readily activated upon oxidation of the nucleotide-free form, we wondered to which extent TRC40 contributes to the protection of cells against H_2_O_2_-induced protein aggregation. We stained control and TRC40-depleted cell with *Proteostat* (Enzo Life Science), a fluorescent probe visualizing stress-induced aggresomes, which are membrane-free cytoplasmic inclusions formed to sequester mis- and unfolded proteins upon proteotoxic stress^44^. As a positive control, we treated cells with MG-132, which inhibits the 26S proteasome and results in massive accumulation of unfolding proteins as shown by a substantial increase in *Proteostat* signal (Figure 4F). We quantified the *Proteostat* signal of individual cells by using the immunofluorescence signal of the cell surface marker integrin-β as internal reference (Figure 4G). Unexpectedly, we found a significantly higher *Proteostat* signal in TRC40-depleted cells compared to the control under non-stress conditions, which might explain the observed growth defect (Figure 4C). Treating cells with 0.5 or 1 mM H_2_O_2_ resulted in a significant increase in the *Proteostat* signal in both, control and TRC40-depleted cells. Yet, the H_2_O_2_-treated control cells were able to efficiently eliminate aggresomes within 60 min, whereas HeLa cells treated with TRC40 siRNA failed to clear the aggregated proteins within the time course of the experiment (Figure 4F and G). In these cells, the *Proteostat* signal remained persistently high. These results demonstrated that TRC40 is crucial to protect cells against acute oxidative stress and plays an essential role in clearing mis- and unfolded proteins, which reveals a so far undiscovered function of TRC40 in restoring proteostasis upon oxidative unfolding stress.

**Figure 4:**
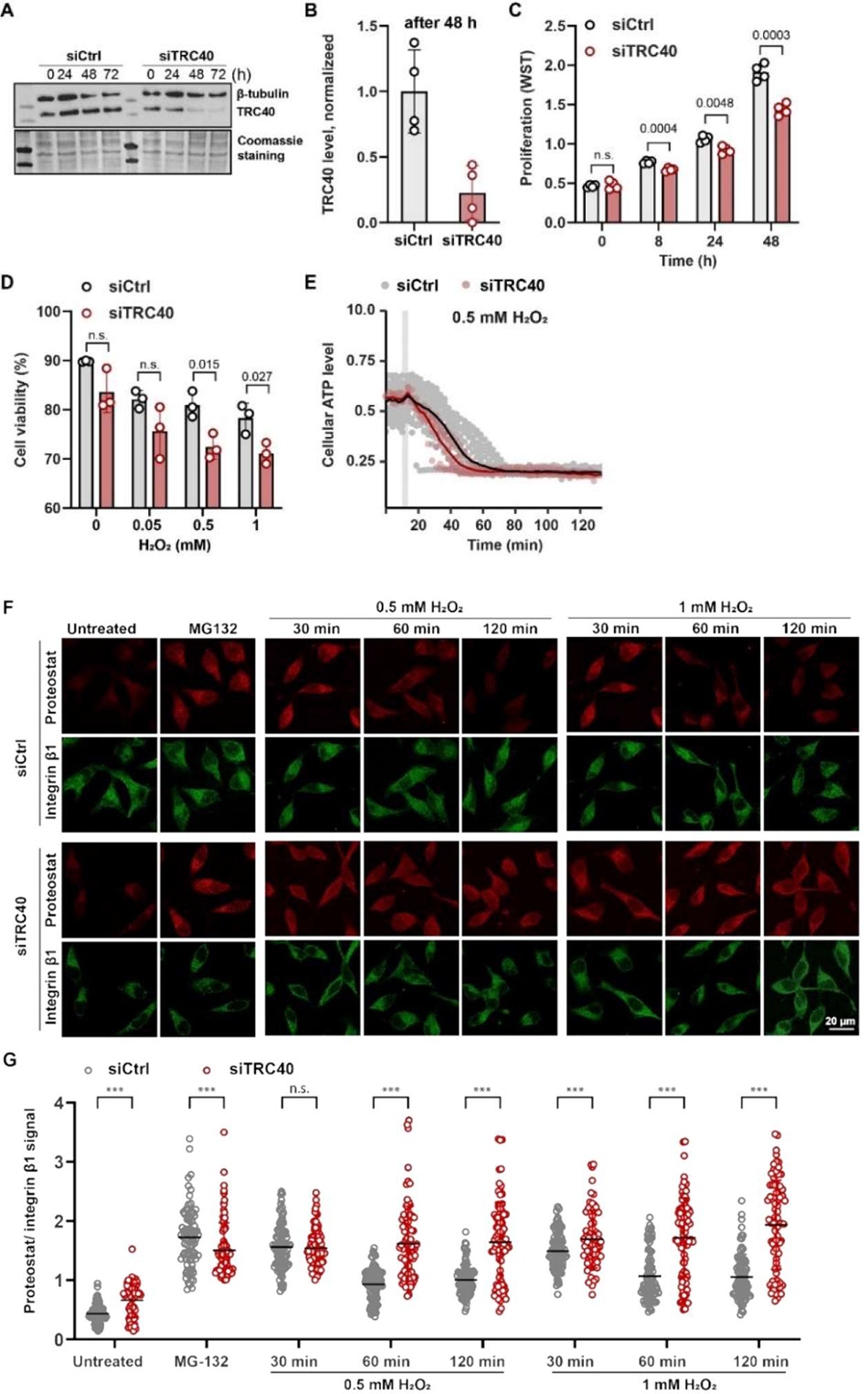
TRC40 protects cellular proteostasis against ATP-depleting oxidative stress. (**A**) HeLa cells were treated with a scrambled siRNA (**siCtrl**) and siRNA targeting TRC40 (**siTRC40**) for up to 72 h. TRC40 protein level were analyzed by Western blotting using an anti-TRC40 antibody. (**B**) TRC40 level 48 h post transfection. (**C**) Cell proliferation analyzed using the WST-1 assay. (**D**) Cells were treated for 30 min with various concentrations of H_2_O_2_ as indicated and cell viability was analyzed by costaining cells with propidium iodide and Annexin V followed by flow cytometry. (**B**-**D**) Data show the mean ± SD of at least three independent experiments. (**E**) Cells were transfected 24 h post RNAi induction with a FRET-based ATP probe. After 48 h, cellular ATP levels were measured for 10 min before and 120 min after adding H_2_O_2_. (**F**) Fluorescent staining of aggresomes using *Proteostat*. Immunofluorescence staining of Integrin β1 was used to visualize the cell surface. As a positive control, cells were treated with the 26S proteasome inhibitor MG-132, inducing accumulation of unfolding proteins in aggresomes. (**G**) The *Proteostat* signal was quantified in comparison to the Integrin β1 signal. Each point represents an individual cell and the data of three independent experiments is shown as mean ± SD. Unpaired, two-sided Student’s t test was applied for statistical analyses and *p*-values < 0.05 are indicated; n.s., not significant

## DISCUSSION

### TRC40 is a multifunctional, redox-regulated proteostasis factor

TRC40 has been well-characterized as ATP-dependent targeting factor for tail-anchored membrane proteins, but certain aspects of its cellular function still remained elusive^25–27,29,32^. Here, we demonstrate an additional facet of TRC40 serving as an ATP-independent chaperone which is essential to cope with protein aggregation upon ATP-depleting, oxidative stress. Chaperone activation is tightly regulated by TRC40’s nucleotide-binding and thiol redox state, a mechanism conserved from its yeast homolog Get3^23^. Nucleotide binding keeps the Get3/TRC40 dimer in a closed and compact conformation, which sterically hinders the accessibly of the thiol groups. Upon nucleotide release, it enters an open conformation which allows the cysteines to readily react with H_2_O_2_ (Figure S2B)^33,34,36^. In clear contrast to yeast Get3, TRC40 is exquisitely sensitive towards oxidation as soon as it is in its nucleotide-free state and readily becomes activated as a molecular chaperone. Nucleotide release is an essential step following ATP hydrolysis and hence, TRC40 always enters a nucleotide-free state during its ATPase cycle, which may favor its oxidation under physiological conditions. However, the millimolar concentration of ATP in the cell^43^ most likely promotes the immediate rebinding of ATP. Yet, thiol oxidation of nucleotide-free TRC40 may occur upon spatiotemporal changes in cellular ATP level, which have been shown to occur in response to metabolic challenges or high energy-consuming processes, such as cell migration, cell division or neuronal signaling^45–47^.

The chaperone function and its regulatory mechanism shed new light on TRC40’s cellular role, which is further supported by the fact that the depletion of TRC40 not only results in reduced cell proliferation, but increases also protein aggregation under normal growth conditions. Notably, for syntaxin 5, which is a *bona fide* tail-anchored protein substrate of TRC40, it has been reported that the property rendering it sensitive towards the loss of TRC40 is mapped to its aggregation-prone, cytoplasmic domain rather than its C-terminal hydrophobic transmembrane anchor^29^. Therefore, it is tempting to speculate that certain proteins depend on TRC40’s chaperone function even under non-stress conditions. Our work supports the notions that the cellular function of TRC40 cannot be reduced to the biogenesis of tail-anchored proteins but needs to be considered as factor of the proteostasis network^29,32^.

### Stress-induced TRC40 foci foster interaction with ATP-dependent chaperones

Based on the switching mechanism, we expected that cellular TRC40 becomes activated under ATP-depleting, oxidative stress conditions. Here, we showed that the exposure of HeLa cells to H_2_O_2_ depletes cellular ATP levels and causes TRC40 to form distinct punctate structures, which rapidly redissolve upon H_2_O_2_ removal. Foci formation has been characterized as a hallmark of chaperone-active Get3 in yeast^21,22^ and serves to sequester mis- and unfolded proteins during stress conditions^48–50^. Unlike toxic amyloid and prion proteins, which are terminally sequestered into insoluble protein deposits (IPODs), stress-damaged proteins are shuttled to dynamic compartments, like aggresomes, Q-bodies and juxtanuclear quality control compartments (JUNQ). To facilitate protein refolding and/or degradation, ATP-dependent chaperones and cochaperones are recruited to these compartments^38^. Our study showed that TRC40 foci contain Hsp70 and are in close proximity to Hsp105/Hsp110. This data is in accordance with studies in yeast showing that Get3 foci colocalize with multiple chaperones, including Ssa2 (Hsp70), Sis1 (Hsp40) and Hsp104^21,22^. Hsp104 is a yeast specific AAA+ disaggregase, well-known for being part of deposition sites of aggregated proteins^51,52,53^. Notably, Hsp104 does not function in isolation, but depends on the Hsp70 system to dissolve protein aggregates^54^. In the metazoan cytosol, a direct Hsp104 homolog does not exist. However, Hsp110 has been demonstrated to cooperate with human Hsp70-Hsp40 to constitute a disaggregase machinery resolubilizing amorphous aggregates^55^. The presence of Hsp70 and Hsp110 near TRC40 foci suggests that TRC40 interacts with ATP-dependent chaperones. While ATP-independent chaperones serve to maintain unfolding protein intermediates in a soluble, folding-competent state, they are not capable of refolding client proteins^13,56^. Instead, these proteins can cooperate with ATP-dependent chaperone systems, like Hsp70/Hsp40, to handover client proteins for refolding^15,16,23^. Although we neither observed an overlap of TRC40 foci with markers of stress granules (G3BP1) nor p-bodies (EDC4), it should be mentioned that Hsp70 is also recruited to these condensates^57^, which may suggest a potential crosstalk among these stress-induced subcellular compartments.

### TRC40 is essential for clearing stress-induced aggresomes

To further elucidate the role of TRC40’s stress-induced chaperone function, we monitored H_2_O_2_-induced formation of aggresomes and aggresome-like inclusions. Aggresomes are dynamic, pericentriolar sequestrations of mis- and unfolded proteins formed upon stress conditions that will impair or overwhelm ATP-dependent chaperones and the ubiquitin-proteasome system^44^. Remarkably, cells were able to cope with these conditions and cleared aggresomes within two hours after adding H_2_O_2_, even though cellular ATP levels have been not restored. Previous studies showed that aggresomes are efficiently cleared via the autophagy-lysosomal pathway in a process known as aggrephagy^58,59^. Consistently, a decrease in cellular ATP level has been shown to induce autophagy via activation of AMP-activated kinase^45^. We made here the surprising discovery that the clearance of H_2_O_2_-induced aggresomes depends on the presence of TRC40 and is profoundly impaired in TRC40-depleted cells, suggesting that TRC40 targets mis- or unfolded proteins to autophagosomal clearance. Notably, a potential link of TRC40 to autophagy has been proposed previously, since overexpression of the cytoplasmic, aggregation-prone domain of syntaxin 5, which strictly depends on TRC40, stimulated autophagy^29^. In how far TRC40 and the formation of TRC40 foci are involved in autophagic clearance of aggregated proteins needs to be now investigated in detail. Our work extends and refines our knowledge about human TRC40, which turned out to be a multifunctional protein protecting proteostasis upon acute oxidative stress conditions and plays a crucial role in the elimination of aggregated proteins.

## ACKNOWLEDGEMENT

We thank the Ulrich, Jakob and Riemer lab as well as Rishav Mitra for helpful discussions. We also thank Robert Keenan, Univeristy of Chicago, for providing us with the plasmid to generate recombinant TRC40. The work was supported by the NRW Rückkehrprogramm of North-Rhine Westphalia, Germany and the internal CRC1218 seed-funding to K.U. and the R35 GM122506 to U.J.. The German Research Foundation (DFG) funds research in the Laboratory of JR through the grants RI2150/5-1 project number 435235019, RI2150/2-2 project number 251546152, RTG2550/1 project number 411422114 and CRC1218 - project number 269925409. B.D. thanks the Sao Paulo Research Foundation (FAPESP), grants #2021/13550-5 and #2019/16224-1.

## AUTHOR CONTRIBUTIONS

**K.U.** Conceptualization; Funding acquisition; Project administration; Investigation; Formal Analysis; Methodology; Supervision; Writing – original draft; Writing – review & editing; **B.D.** Conceptualization; Investigation; Formal Analysis; Methodology; **O.C.** Investigation; **R.D.** Investigation; **K.L.** Formal Analysis **J.R.** Formal Analysis; **U.J.** Conceptualization; Funding acquisition; Writing – review & editing

## MATERIAL AND METHODS

### RESOURCE AVAILABILITY

#### Lead Contact

Further information and requests for resources, reagents and plasmids should be directed to and will be fulfilled by the corresponding author, Kathrin Ulrich.

## METHOD DETAILS

### Expression and purification of recombinant protein

*E. coli* BL21 DE3 cells, transformed with the pET28-6xHis-Tev-DrTRC40 plasmid for IPTG-inducible expression of TRC40 from *Danio rerio*, were inoculated in 100 ml of LB containing 50 μg/ml kanamycin (LB-Kan) and incubated while shaking at 160 rpm overnight at 37°C. The overnight culture was diluted 1:70 in 4 x 1 l LB-Kan and incubated while shaking at 37°C until an OD_600_ of 0.5 was reached. Protein expression was induced by adding 0.4 mM IPTG and the culture was incubated for 4 h at 30°C. Cells were pelleted and stored at −80°C. Cell pellets were resuspended in extraction buffer (50 mM Tris, 50 mM NaCl, 2 mM magnesium acetate, 1 mM imidazole, pH 7.5), supplemented with one tablet of EDTA-free protease inhibitor (complete EDTA-free Protease Inhibitor Cocktail, Roche), 1 mM PMSF, 2 mg/ml lysozyme, 2 mM DTT and a spoon tip DNase. Cells were lysed by sonication on ice and the crude extract was centrifuged at 36,000 x g for 45 min at 4°C. The supernatant was filtered through a 0.45 µm filter and loaded onto a HisTrap HP column. The column was washed with 5 column volumes (CVs) of extraction buffer, 2 CVs of 4 mM ATP in extraction buffer, 2 CVs of 450 mM NaCl in extraction buffer, and 2 CVs of extraction buffer until the flow-through was protein-free. Next, bound proteins were eluted with 500 mM imidazole in the extraction buffer. The protein-containing fractions were collected and analyzed on a reducing NuPAGE Bis-Tris protein gel (Thermo Scientific) stained with Coomassie blue. The fractions containing the TRC40-fusion protein were pooled and the His-tag was cleaved by adding TEV protease during dialysis in 3 l of dialysis buffer (50 mM Tris, 40 mM NaCl, 5 mM magnesium acetate, 0.5 mM DTT, pH 7.5) overnight at 4°C. The cleaved protein was applied again to the HisTrap column and the flow-through containing tag-free TRC40 was collected. The cleavage efficiency and purity of the purified protein were tested on a reducing NuPAGE Bis-Tris protein gel stained with Coomassie. The protein concentration was determined spectroscopically at 280 nm using an extinction coefficient of 14.44 mM^-1^ cm^-1^ (assuming that all cysteines are reduced). The purification of recombinant His-tagged Get3 from *Saccharomyces cerevisiae* was performed as described in Ulrich et al. 2022.

### TRC40/Get3 reduction, oxidation, re-reduction, and thiol blocking

To generate fully reduced TRC40 or Get3, the recombinant protein was diluted to 5 µM in 40 mM HEPES-KOH, pH 7.5, 5 mM DTT, 2 mM Mg-ATP and 5 µM ZnCl_2_, and incubated overnight at 4°C with gentle shaking. The mixture was transferred to an Amicon-Ultra centrifugal filter with a 30 kDa cut-off to concentrate the protein to about 80-100 µM and washed four times with 15 ml HEPES-KOH, pH 7.5 to completely remove DTT. Finally, the protein was run through a gel filtration spin column (Zeba Spin Desalting Column, Thermo Scientific) to completely remove any unbound ATP. To generate nucleotide-free TRC40/Get3, 50 µM of protein was incubated for 30 min at 30°C with 5 U apyrase in 100 µl HEPES-KOH, pH 7.4.

For oxidation, 50 µM TRC40/Get3 was treated with 2 mM H_2_O_2_ and 50 µM CuCl_2_ for 10 min at 37°C. To study milder oxidizing conditions, the H_2_O_2_/CuCl_2_ mixture was replaced by 5 mM H_2_O_2_. For the re-reduction, the oxidized protein was diluted to 5 µM in 40 mM HEPES-KOH, pH 7.5, and incubated in the presence of 5 mM DTT, 5 µM ZnCl_2_, and 2 mM Mg-ATP for 4 h at 30°C. To irreversibly block thiol residues, 50 µM TRC40 was incubated in 40 mM HEPES-KOH, pH 7.5 with 10 mM *N*-ethylmaleimide (NEM) for 30 min at 25°C in the dark.

### Chaperone activity assay

The chaperone activity was analyzed using the citrate synthase aggregation assay ^60,61^. Citrate synthase (12 µM) was chemically denatured in 40 mM HEPES-KOH, pH 7.5 containing 6 M guanidine hydrochloride overnight at RT. To follow protein aggregation, citrate synthase was diluted to a final concentration of 0.15 µM in 1.6 ml 40 mM HEPES-KOH, pH 7.5 at 30°C in a quarz cuvette under continuous stirring. Light scattering was monitored (λ_ex/em_ = 360 nm) using a Hitachi F4500 fluorescence spectrophotometer equipped with a temperature-controlled cuvette holder and stirrer. After 4 min of incubation, the signal reached a plateau, which was set to 0% chaperone activity (completely aggregated protein). To test the effects of TRC40 or Get3, the buffer was supplemented with a 4-fold molar excess of total protein over citrate synthase before citrate synthase was added. The light scattering detected in the presence of TRC40 oxidized by 2 mM H_2_O_2_/50 mM CuCl_2_ for 10 min at 37°C was set to 100% chaperone activity for normalization.

### ATPase activity assay

The ATPase activity was measured using an NADH-coupled enzymatic assay containing 1 mM phosphoenolpyruvate, 0.5 mM NADH, 12 U pyruvate kinase, and 12 U lactate dehydrogenase in 100 mM HEPES-KOH, 100 mM KCl, 10 mM MgCl_2_, 20% glycerol, pH 7.5 at 30°C ^22,62^. After pre-incubating 8 µM TRC40 in the assay buffer for 5 min, 2 mM Mg-ATP was added to start the reaction. Upon hydrolysis of ATP to ADP, pyruvate kinase catalyzes the transfer of a phosphate group from phosphoenolpyruvate, yielding one molecule of pyruvate and regenerating one molecule of ATP. Lactate dehydrogenase catalyzes the reduction of pyruvate to lactate under the consumption of one molecule NADH per one molecule of pyruvate. The linear decrease of NADH was measured over time at 340 nm (ε_340 nm_ = 6.2 mM^-1^ cm^-1^) in a 96-well BMG FLUOstar Omega microplate reader. Using Lambert-Beer’s law, the amount of hydrolyzed ATP per minute (µmol/min = U) in the assay volume was calculated and used to determine the specific enzyme ATPase activity (U/µg).

### Analytical gel filtration

The oligomeric state of TRC40 and Get3 were monitored by size-exclusion chromatography using a Superdex 200 10/300 GL column (GE Healthcare) equilibrated with 40 mM HEPES, 140 mM NaCl, pH 7.5 as described previously ^22,23^. Briefly, 200 µl of 50 µM protein was loaded onto the column and eluted with a flow rate of 0.5 ml/min at 4°C using an Akta-FPLC system (GE Healthcare). Absorption at 280 nm was used to monitor the protein elution.

### Reverse thiol labeling with mPEG-MAL

TRC40 was mixed with 20 mM NEM in denaturing alkylating buffer (DAB, 200 mM Tris-HCl, 10 mM EDTA, 6 M urea, 0.5% w/v SDS, pH 8.5) ^63^ and incubated for 30 min in the dark to irreversibly block all reduced cysteine residues. The protein was precipitated by adding 10% v/v ice-cold trichloroacetic acid (TCA). After 45 min on ice, samples were centrifuged for 10 min at 16,000 x g, 4°C and washed once with 10% v/v and 5% v/v ice-cold TCA to remove excess NEM. The pellet was immediately re-dissolved in 40 µl DAB containing 2 mM DTT. After 30 min at 37 °C, 10 µl of 50 mM methoxy polyethylene glycol maleimide (mPEG-MAL 5,000 Da) was added and samples were incubated for 1 h at RT. Samples were mixed with SDS-sample buffer and boiled for 5 min. Excess mPEG-MAL was quenched by adding 50 mM DTT and samples were loaded onto a 4-12% NuPAGE Bis-Tris gels followed by Coomassie staining.

### Cell culture

HeLa (EM-2-11ht) cells were cultured at 5% CO_2_ at 37°C in DMEM (Life Technologies) supplemented with 10% fetal bovine serum and 1% penicillin and streptomycin. Cells were split every two days until 15 passages were reached. Therefore, cells were treated with 0.25% trypsin for 5 min at 37°C, resuspended in fresh DMEM and proceeded to cell counting. For each experiment, cells were seeded two days prior the treatment to a density which did not exceed 60-80% confluceny on the day of the experiment.

### Immunofluorescence staining of TRC40

HeLa cells were seeded (5 x 10^4^ cells/well) onto 12 mm glass coverslips placed in 24-well plates. After 48 h, cells were washed with PBS and incubated in HBSS buffer with or without 0.5 mM H_2_O_2_. After 15 min, cells were fixed with 4% paraformaldehyde in PBS for 20 min, permeabilized with 0.3% Triton X-100 in PBS and blocked with 2% BSA in PBS for 1 h with PBS washes between the individual steps. Coverslips were incubated with primary antibodies diluted (1:1,000) in 2% BSA overnight at RT. After incubation, cells were washed four times with PBS and incubated with secondary antibodies for 2 h at RT. Cells were washed three times with PBS and incubated with 2.5 μg/ml DAPI for 15 min. After two more PBS washes, coverslips were mounted on microscope objective slides using ProLongGold Antifade Reagent. Images were acquired with a Leica SP8 laser scanning confocal microscope (Leica GmbH, Mannheim, Germany) on a DMI8 microscope base using LAS X software, a 63x oil objective, and a 405 nm diode laser, in addition to a multi-line white light laser, set to an excitation wavelength of 488 nm and 594 nm. Spectral detection using a PMT from 410 to 450 nm was utilized for DAPI, a HyD detector from 490 to 560 nm for AlexaFluor 488, and a HyD detector from 604 to 640 nm for AlexaFluor 594.

### Transient transfection of HeLa cells

HeLa cells (1×10^6^ cells/well) were seeded in 6-well plates and incubated for 24 h. RNA-lipofectamine RNAiMAX mix was prepared following the manufacturer’s instructions for control siRNA (Silencer Negative Control No. 1 siRNA, AM4611) and TRC40 siRNA (s1675) as reported previously ^39^, yielding a final concentration of 50 nM in Opti-MEM medium. The mixture was incubated for 5 min and added to cells, containing DMEM-F12 medium. After 24 h, the cell medium was changed and cells were incubated for another 24 h before the experiments.

### Protein quantification from cell lysates

Protein concentration of human serum was determined using Pierce Bicinchoninic Acid (BCA) Protein Assay Kit (Thermo Fisher Scientific) with bovine albumin as standard following the manufacturer’s instructions.

### Western blot analysis

After transfection, cells were washed twice with PBS and scrapped in lysis buffer (RIPA buffer, 1 mM PMSF, protease inhibitor cocktail, and 1 mM EDTA). Samples were incubated for 30 min on ice and spun down (4 °C, 20 min, 12,000 rpm). The soluble supernatant was collected and the total protein concentration was determined. Samples were mixed with SDS-sample buffer and boiled for 5 min. A total amount of 20 µg per sample was applied onto a 4-12% NuPAGE Bis-Tris gels. Proteins were transferred to a PVDF membrane using the Trans-Blot Turbo Transfer System (Bio-Rad). The membrane was blocked for 1 h with 5% milk in PBS and incubated with primary antibodies against TRC40 (rabbit, 1:1000 in 5% milk in TBS-T) and β-tubulin (mouse, 1:1000 in 5% milk in TBS-T) overnight at 4 °C. After washing the membrane three times with PBS, IRDye 800CW anti-rabbit and IRDye 680RD anti-mouse secondary antibodies were diluted 1:5,000 in 5% milk/PBS and added to the membrane for 1 h at RT. After washing three times with PBS, fluorescent signals were detected using the Odyssey CLx scanner (LI-COR).

### WST proliferation assay

HeLa cells were transfected as described above. After 24 h, cells were seeded onto 96-well cell culture plates (10^4^ cells/well) and the WST-1 Cell Proliferation Assay kit was used to assess proliferation after 8 h, 24 h, and 48 h. Briefly, the cell medium was removed, cells were washed with PBS. The mixture of the WST-1 Developer Reagent and the Electron Mediator Solution was first diluted in fresh DMEM to the working concentration following the manufacturer’s instruction and then added to the cells. Absorption was measured in a Tecan Infinite M1000 microplate reader at a wavelength of 450 nm.

### Cell viability analysis using flow cytometry

HeLa cells were transfected as described above. After 48 h, cells were harvested, resuspended in HBSS buffer (10^6^ cells/ml) and incubated without and with various concentrations of H_2_O_2_ for 2 h at 37°C with agitation at 300 rpm. Afterwards, cells were centrifuged and stained with Annexin V-FITC Apoptosis Detection Kit (APOAF-60TST, Sigma Aldrich) according to the manufacturer’s instructions, and subsequently subjected to flow cytometry. Samples were analyzed using a CytoFLEX SRT flow cytometer (Beckman Coulter). Fluorescence emitted by Annexin V-FITC and PI was measured at 525 nm and 690 nm, respectively, upon excitation at 488 nm. Single-stained compensation controls were used to calculate the compensation matrix. 30,000 events were acquired and recorded per sample. Data were analyzed using FlowJo version 10 software.

### Proteostat aggregation assay

The *Proteostat* Aggresome detection kit (Enzo Life Sciences) was used to visualize mis- and unfolded proteins. Cells were 24 h post transfection seeded onto a 24-well plate (2×10^4^ cells/well) containing 12 mm glass coverslips. After 24 h, cells were washed with PBS and treated with 0.5 or 1 mM H_2_O_2_ in HBSS buffer at 37 °C. After incubation of 30, 60, and 120 min, cells were fixed with 4% paraformaldehyde for 20 min, following the immunofluorescence protocol described before. After permeabilization and blocking, coverslips were incubated with integrin β1 primary antibody overnight at RT. After washing with PBS, coverslips were incubated with a secondary antibody conjugated with Alexa Fluor 488 for 2 h at RT. Cells were washed twice with PBS and the *Proteostat* dye was added at 1:4,000 dilution in PBS for 30 min at RT. After washing twice with PBS, cells were mounted and imaged using a Leica SP8 laser scanning confocal microscope (Leica) on a DMI8 microscope base using LAS X software, a 63× oil objective and a 405 nm diode laser, in addition to a multi-line white light laser set to 488 and 594 nm excitation wavelengths with default acquiring configuration for Rhodamine (for *Proteostat*) and FITIC (for integrin β1). Fluorescence intensity was quantified using Image J software. *Proteostat* signal of each individual cell was normalized by the respective integrin β1 signal.

### Data analysis

All experiments, if not otherwise indicated, were conducted with n equal or higher than three. Error bars represent the average ± standard deviation of the mean. Statistical significance was determined using the unpaired, one-sided Student’s t-test.

## SUPPLEMENT

**Figure S1:**
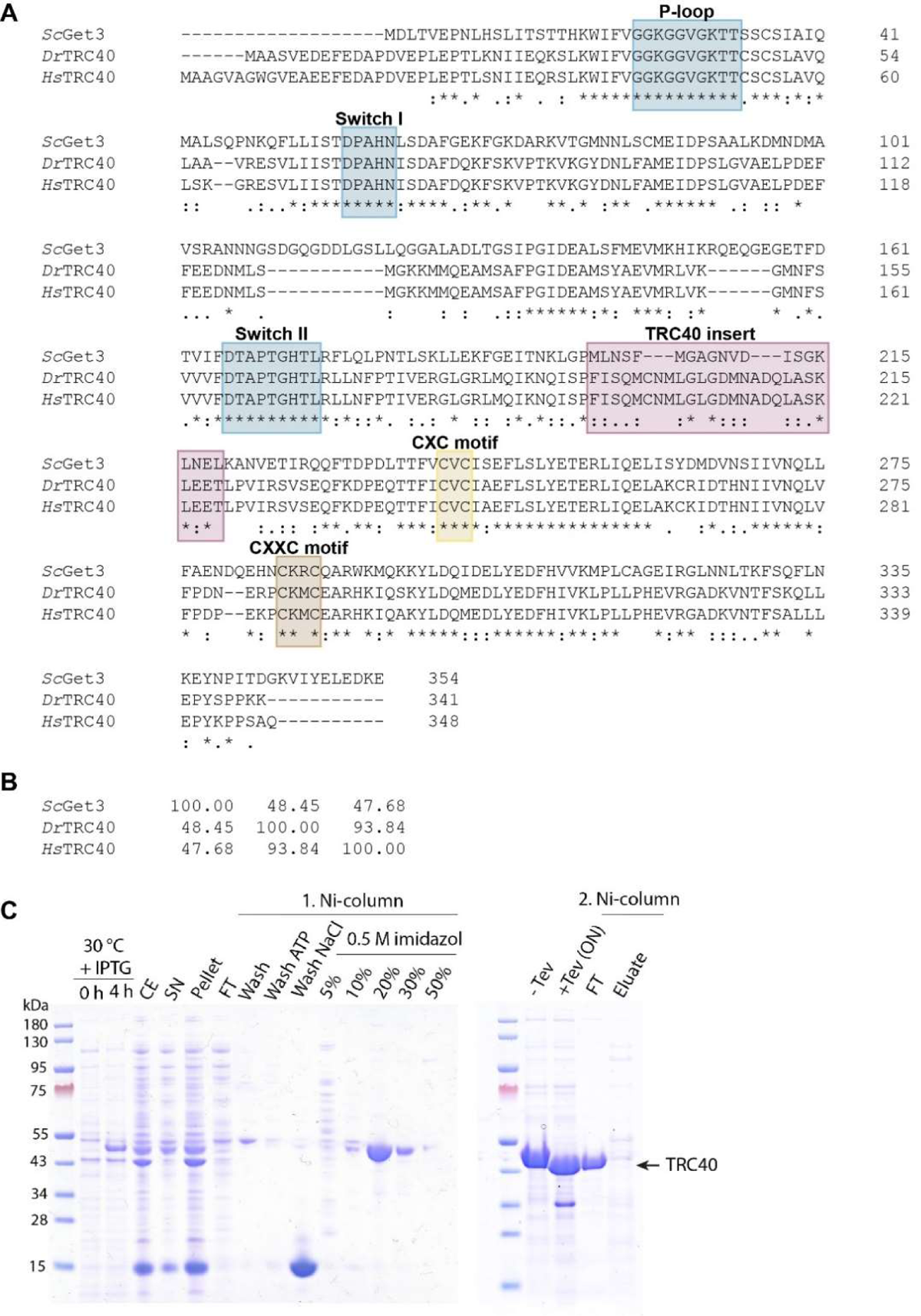
A) Sequence alignment of Get3 from *Saccharomyces cerevisiae* (*UniProt ID*: Q12154) and TRC40 from *Danio rerio* (*UniProt ID*: Q6IQE5) and *Homo sapiens* (*UniProt ID*: O43681). Conserved structural elements are highlighted. B) Percent identity matrix. The alignment was generated with *Clustal Omega* v 1.2.4. C) Expression and purification of recombinant, His-tagged *Dr*TRC40 as described in Material and Methods. Digest with the Tev protease cleaved the His-tag, resulting in pure and soluble full-length TRC40. CE, crude extract; SN, supernatant; FT, flow-through

**Figure S2:**
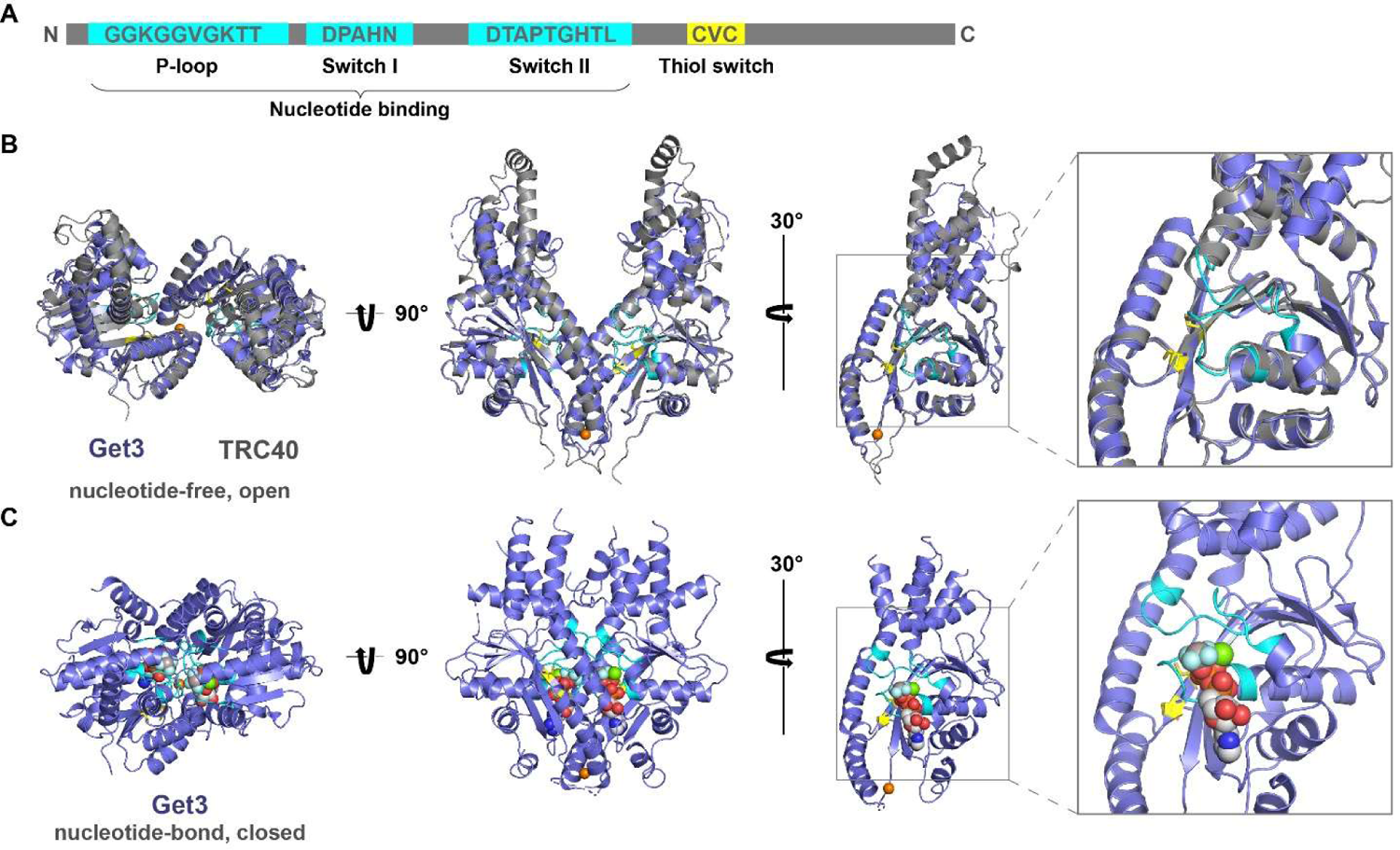
Structural conservation and nucleotide-dependent conformational changes of Get3 A) Schematic representation of the nucleotide-binding domain (cyan) and the CXC-motif (yellow) conserved in Get3 and TRC40. B) Alignment of the Get3 homodimer (in purple) in its nucleotide-free state (PDB ID 2WOO, Mateja et al. 2009) and the Alphafold structure of human TRC40 (in grey). C) In the presence of ATP, Get3’s helical subdomains undergo conformational rearrangements and become intimately associated which turns Get3 into its closed conformation. Get3 is shown in its Mg^2+^-ADP-AlF_4_^-^ -bound state, which mimic the ATP-bound state (PDB ID 2WOI, Mateja et al. 2009). The insets show a zoomed-in view into the nucleotide-binding site and the CXC-motif of the monomeric subunit.

**Figure S3:**
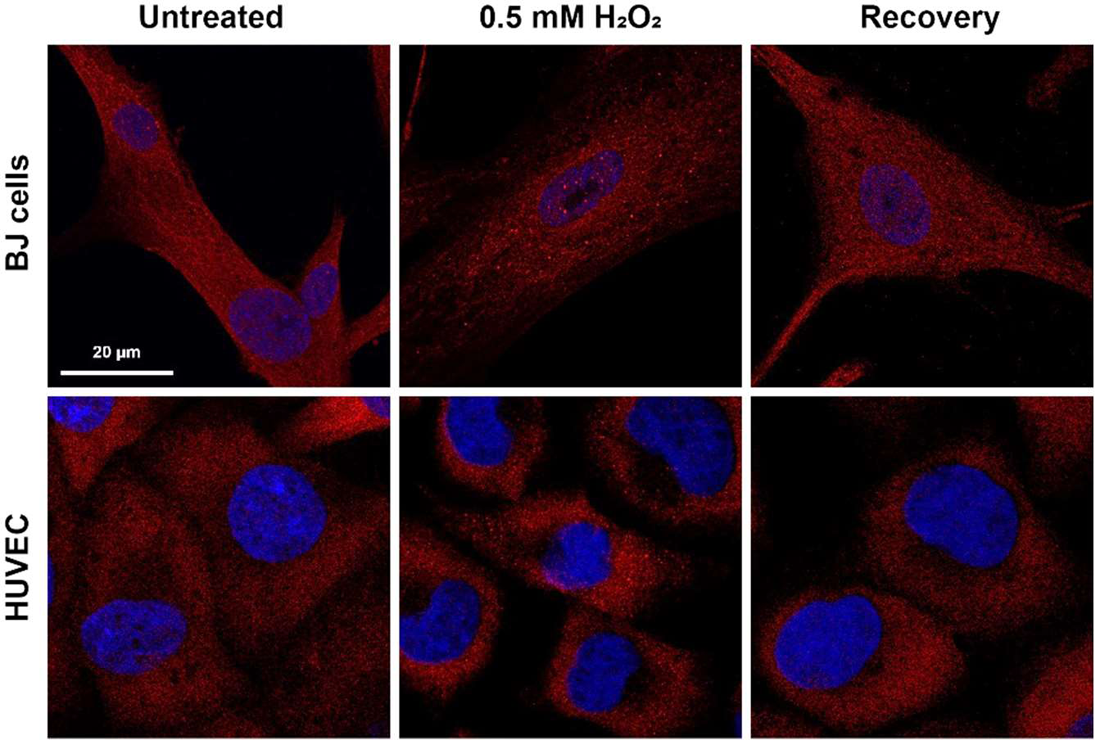
H_2_O_2_-induced foci in human fibroblasts (BJ cells) and human umbilical vein endothelial cells (HUVEC) Cells were treated with 0.5 mM H_2_O_2_ in HBSS buffer for 15 min. After washing twice, cells were supplemented with fresh DMEM. Human TRC40 was analyzed by immunofluorescence staining using an anti-TRC40 antibody.

---

**Table 1:** Resources.

| Reagent or Resource | Source | Identifier |
| --- | --- | --- |
| **Antibodies** |  |  |
| Anti-ASNA1/TRC40, rabbit polyclonal | Thermo Fisher Scientific | Cat# PA576471; RRID AB_2720198 |
| Anit-Beta-tubulin, mouse monoclonal | Thermo Fisher Scientific | Cat# 32-2600; RRID AB_2533072 |
| Anti-Rabbit IgG Alexa Fluor 594, goat polyclonal | Thermo Fisher Scientific | Cat# A11012; RRID AB_2534079 |
| Anti-Mouse IgG Alexa Fluor 488, goat polyclonal | Thermo Fisher Scientific | Cat# A11001; RRID AB_2534069 |
| Anti-HSP70, mouse monoclonal | Santa Cruz Biotechnology | Cat# sc-32239; RRID AB_627759 |
| Anti-HSP105, mouse monoclonal | Santa Cruz Biotechnology | Cat# sc-74550; RRID AB_2119247 |
| Anti-Edc4, mouse monoclonal | Santa Cruz Biotechnology | Cat# sc-376382; RRID AB_10988077 |
| Anti-SQSTM1/p62, rabbit polyclonal | Cell Signaling Technology | Cat# 5114; RRID AB_10624872 |
| Anti-G3BP1, mouse monoclonal | BD Biosciences | Cat# 611126; RRID AB_398437 |
| Anti-Integrin beta 1, mouse monoclonal | Abcam | Cat# ab30394; RRID AB_775726 |
| IRDye Anti-Rabbit, 800CW, donkey polyclonal | LI-COR | 925-32211 |
| IRDye Anti-Mouse, 680RD, donkey polyclonal | LI-COR | 925-68070 |
| **Bacterial strains** |  |  |
| *E. coli* BL21 (DE3) | Novagen | Cat#69450 |
| **Cell culture components** |  |  |
| DMEM, high glucose, pyruvate | Gibco, Thermo Fisher Scientific | Cat# 11-995-065 |
| Penicillin-streptomycin 100x | Gibco, Thermo Fisher Scientific | Cat# 15140-122 |
| Phosphate Buffered Saline (PBS) | Gibco, Thermo Fisher Scientific | Cat# 10010031 |
| Hanks' Balanced Salt Solution (HBSS) | Gibco, Thermo Fisher Scientific | Cat# 14025092 |
| Trypsin-EDTA 0.25% | Gibco, Thermo Fisher Scientific | Cat# 25200056 |
| Opti-MEM Reduced Serum Medium | Gibco, Thermo Fisher Scientific | Cat# 31985088 |
| DMEM/F-12 | Gibco, Thermo Fisher Scientific | Cat# 11320033 |
| Fetal Bovine Serum (FBS) | Sigma-Aldrich | Cat# F4135-500ML |
| Lipofectamine RNAiMAX Transfection Reagent | Invitrogen, Thermo Fisher Scientific | Cat# 13778075 |
| **Chemicals and recombinant proteins** |  |  |
| Apyrase | NEB | Cat#M0398L |
| Bovine Serum Albumin (BSA) | Sigma-Aldrich | A7906 |
| Citrate synthase from porcine heart | Sigma-Aldrich | Cat#C3260 |
| 1,4-Dithiothreitol (DTT) | Sigma-Aldrich | Cat# 10708984001 |
| Dnase | Thermo Fisher Scientific | Cat#EN0521 |
| EDTA-free protease inhibitor cocktail | Sigma-Aldrich | Cat#11873580001 |
| Hydrogen peroxide (H<sub>2</sub>O<sub>2</sub>), 30% | Thermo Fisher Scientific | Cat#H325 |
| Kanmycin monosulfate | Gold Biotechnology | Cat# K-120-5 |
| Lactate dehydrogenase | Roche | Cat#10127230001 |
| Lysozyme | MP Biomedicals | Cat# 0210083110 |
| Methoxypolyethylene glycol maleimide (mPEG-MAL), 5,000 Da | Sigma-Aldrich | Cat#63187 |
| NADH | Sigma-Aldrich | Cat#N8129 |
| N-ethylmaleimide (NEM) | Sigma-Aldrich | Cat#E3876 |
| Paraformaldehyde, 8% | Electron Microscopy Sciences | Cat# 1578100 |
| Phenylmethylsulfonyl fluoride (PMSF) | Sigma-Aldrich | Cat #10837091001 |
| Phosphoenolpyruvate | Sigma-Aldrich | Cat#860077 |
| ProLongGold Antifade Reagent | Cell Signaling | Cat# 9071 |
| Pyruvate kinase | Sigma-Aldrich | Cat#P1506 |
| TEV protease | Produced in-house | N/A |
| Triton X-100 | Sigma-Aldrich | Cat# T9284 |
| **Commercial assay kits** |  |  |
| Bicinchoninic Acid (BCA) Protein Assay Kit, Pierce | Thermo Fisher Scientific | Cat# A55860 |
| WST-1 Cell Proliferation Assay Kit | Cayman Chemical | Cat#10008883 |
| *Proteostat* Aggresome detection kit | Enzo Life Sciences | ENZ-51035 |
| Annexin V-FITC Apoptosis Detection kit | Sigma-Aldrich | Cat# APOAF-60TST |
| **Materials** |  |  |
| Amicon Ultra-15, Centrifugal filter, 30 kDa cut-off | Thermo Fisher Scientific | Cat#UFC903096 |
| Dialysis membrane tubing, MWCO 6000-8000 | Thermo Fisher Scientific | Cat#08-670A |
| HisTrap High Performance (HP) column, 5 ml | GE Healthcare/Cytiva | Cat#17-5248-02 |
| HiTrap Q HP column, 5 ml | GE Healthcare/Cytiva | Cat#17-1154-01 |
| NuPAGE 4-12% Bis-Tris polyacrylamide gels | Thermo Fisher Scientific | Cat#NP0322BOX |
| Superdex 200 10/300 GL column | Sigma-Aldrich | Cat#GE-17-5175-01 |
| Zeba Spin Desalting columns, 2 ml | Thermo Fisher Scientific | Cat#89890 |
| **Oligonucleotides** |  |  |
| Silencer Select ASNA1 ID s1675 | Thermo Fisher Scientific | Cat#4390824 |
| Silencer Negative Control No.1 siRNA | Thermo Fisher Scientific | Cat#AM4611 |
| **Plasmids** |  |  |
| pcDNA-AT1.03 | Imamura et al. 2009 | N/A |
| pET28-6His-DrTRC40 | Mariappan et al. 2011 | N/A |
| pQE80-10His-ZZ-scGet3, wild-type | Voth et al. 2014 | N/A |
| **Software** |  |  |
| Adobe Illustrator 2024 | Adobe | https://www.adobe.com/adobe/illustrator |
| Fiji Image J 1.53q | Image J | imagej.nih.gov/ij |
| FlowJo v10.8.1 | BD Biosciences | https://www.flowjo.com/ |
| GraphPad Prism 10.1.2 | GraphPad | https://www.graphpad.com/ |
| ImageStudioLite 2.1 | LI-COR | https://www.licor.com/bio/imagestudio-lite/ |
| Leica Application Suite X 3.7.6.25997 | Leica | https://www.leica-microsystems.com/products/microscope-software |

## REFERENCES

1. Hartl, F.U., Bracher, A., and Hayer-Hartl, M. (2011). Molecular chaperones in protein folding and proteostasis. Nature 475, 324–332. 10.1038/nature10317.

2. Kim, Y.E., Hipp, M.S., Bracher, A., Hayer-Hartl, M., and Hartl, F.U. (2013). Molecular chaperone functions in protein folding and proteostasis. Annu Rev Biochem 82, 323–355. 10.1146/annurev-biochem-060208-092442.

3. Rosenzweig, R., Nillegoda, N.B., Mayer, M.P., and Bukau, B. (2019). The Hsp70 chaperone network. Nat Rev Mol Cell Biol 20, 665–680. 10.1038/s41580-019-0133-3.

4. Mitra, R., Wu, K., Lee, C., and Bardwell, J.C.A. (2022). ATP-Independent Chaperones. Annu Rev Biophys 51, 409–429. 10.1146/annurev-biophys-090121-082906.

5. Ulrich, K., and Jakob, U. (2019). The role of thiols in antioxidant systems. Free Radic Biol Med 140, 14–27. 10.1016/j.freeradbiomed.2019.05.035.

6. Reichmann, D., Voth, W., and Jakob, U. (2018). Maintaining a healthy proteome during oxidative stress. Mol Cell 69, 203–213. 10.1016/j.molcel.2017.12.021.

7. Winter, J., Linke, K., Jatzek, A., and Jakob, U. (2005). Severe oxidative stress causes inactivation of DnaK and activation of the redox-regulated chaperone Hsp33. Mol Cell 17, 381–392. 10.1016/j.molcel.2004.12.027.

8. Ralser, M., Wamelink, M.M., Kowald, A., Gerisch, B., Heeren, G., Struys, E.A., Klipp, E., Jakobs, C., Breitenbach, M., Lehrach, H., and Krobitsch, S. (2007). Dynamic rerouting of the carbohydrate flux is key to counteracting oxidative stress. J Biol 6, 10. 10.1186/jbiol61.

9. Kumsta, C., Thamsen, M., and Jakob, U. (2011). Effects of oxidative stress on behavior, physiology, and the redox thiol proteome of *Caenorhabditis elegans*. Antioxid Redox Signal 14, 1023–1037. 10.1089/ars.2010.3203.

10. Ulrich, K. (2023). Redox-regulated chaperones in cell stress responses. Biochem Soc Trans 51, 1169–1177. 10.1042/BST20221304.

11. Goemans, C.V., and Collet, J.F. (2019). Stress-induced chaperones: a first line of defense against the powerful oxidant hypochlorous acid. F1000Res *8*. 10.12688/f1000research.19517.1.

12. Muller, A., Langklotz, S., Lupilova, N., Kuhlmann, K., Bandow, J.E., and Leichert, L.I. (2014). Activation of RidA chaperone function by N-chlorination. Nat Commun 5, 5804. 10.1038/ncomms6804.

13. Jakob, U., Muse, W., Eser, M., and Bardwell, J.C. (1999). Chaperone activity with a redox switch. Cell 96, 341–352. 10.1016/s0092-8674(00)80547-4.

14. Becker, S.H., Ulrich, K., Dhabaria, A., Ueberheide, B., Beavers, W., Skaar, E.P., Iyer, L.M., Aravind, L., Jakob, U., and Darwin, K.H. (2020). *Mycobacterium tuberculosis* Rv0991c is a redox-regulated molecular chaperone. mBio 11. 10.1128/mBio.01545-20.

15. Teixeira, F., Castro, H., Cruz, T., Tse, E., Koldewey, P., Southworth, D.R., Tomas, A.M., and Jakob, U. (2015). Mitochondrial peroxiredoxin functions as crucial chaperone reservoir in *Leishmania infantum*. Proc Natl Acad Sci U S A 112, E616–624. 10.1073/pnas.1419682112.

16. Currier, R.B., Ulrich, K., Leroux, A.E., Dirdjaja, N., Deambrosi, M., Bonilla, M., Ahmed, Y.L., Adrian, L., Antelmann, H., Jakob, U., et al. (2019). An essential thioredoxin-type protein of *Trypanosoma brucei* acts as redox-regulated mitochondrial chaperone. PLoS Pathog 15, e1008065. 10.1371/journal.ppat.1008065.

17. Aramin, S., Fassler, R., Chikne, V., Goldenberg, M., Arian, T., Kolet Eliaz, L., Rimon, O., Ram, O., Michaeli, S., and Reichmann, D. (2020). TrypOx, a novel eukaryotic homolog of the redox-regulated chaperone Hsp33 in *Trypanosoma brucei*. Front Microbiol 11, 1844. 10.3389/fmicb.2020.01844.

18. Goemans, C.V., Vertommen, D., Agrebi, R., and Collet, J.F. (2018). CnoX is a chaperedoxin: a holdase that protects its substrates from irreversible oxidation. Mol Cell 70, 614–627 e617. 10.1016/j.molcel.2018.04.002.

19. Schuldiner, M., Metz, J., Schmid, V., Denic, V., Rakwalska, M., Schmitt, H.D., Schwappach, B., and Weissman, J.S. (2008). The GET complex mediates insertion of tail-anchored proteins into the ER membrane. Cell 134, 634–645. 10.1016/j.cell.2008.06.025.

20. Stefanovic, S., and Hegde, R.S. (2007). Identification of a targeting factor for posttranslational membrane protein insertion into the ER. Cell 128, 1147–1159. 10.1016/j.cell.2007.01.036.

21. Powis, K., Schrul, B., Tienson, H., Gostimskaya, I., Breker, M., High, S., Schuldiner, M., Jakob, U., and Schwappach, B. (2013). Get3 is a holdase chaperone and moves to deposition sites for aggregated proteins when membrane targeting is blocked. J Cell Sci 126, 473–483. 10.1242/jcs.112151.

22. Voth, W., Schick, M., Gates, S., Li, S., Vilardi, F., Gostimskaya, I., Southworth, D.R., Schwappach, B., and Jakob, U. (2014). The protein targeting factor Get3 functions as ATP-independent chaperone under oxidative stress conditions. Mol Cell 56, 116–127. 10.1016/j.molcel.2014.08.017.

23. Ulrich, K., Farkas, A., Chan, O., Katamanin, O., Schwappach, B., and Jakob, U. (2022). From guide to guard-activation mechanism of the stress-sensing chaperone Get3. Mol Cell 82, 3226–3238 e3227. 10.1016/j.molcel.2022.06.015.

24. Farkas, A., De Laurentiis, E.I., and Schwappach, B. (2019). The natural history of Get3-like chaperones. Traffic 20, 311–324. 10.1111/tra.12643.

25. Borgese, N., Coy-Vergara, J., Colombo, S.F., and Schwappach, B. (2019). The ways of tails: the GET pathway and more. Protein J 38, 289–305. 10.1007/s10930-019-09845-4.

26. Mariappan, M., Mateja, A., Dobosz, M., Bove, E., Hegde, R.S., and Keenan, R.J. (2011). The mechanism of membrane-associated steps in tail-anchored protein insertion. Nature 477, 61–66. 10.1038/nature10362.

27. Favaloro, V., Spasic, M., Schwappach, B., and Dobberstein, B. (2008). Distinct targeting pathways for the membrane insertion of tail-anchored (TA) proteins. J Cell Sci 121, 1832–1840. 10.1242/jcs.020321.

28. Mukhopadhyay, R., Ho, Y.S., Swiatek, P.J., Rosen, B.P., and Bhattacharjee, H. (2006). Targeted disruption of the mouse Asna1 gene results in embryonic lethality. FEBS Lett 580, 3889–3894. 10.1016/j.febslet.2006.06.017.

29. Rivera-Monroy, J., Musiol, L., Unthan-Fechner, K., Farkas, A., Clancy, A., Coy-Vergara, J., Weill, U., Gockel, S., Lin, S.Y., Corey, D.P., et al. (2016). Mice lacking WRB reveal differential biogenesis requirements of tail-anchored proteins *in vivo*. Sci Rep 6, 39464. 10.1038/srep39464.

30. Mateja, A., Szlachcic, A., Downing, M.E., Dobosz, M., Mariappan, M., Hegde, R.S., and Keenan, R.J. (2009). The structural basis of tail-anchored membrane protein recognition by Get3. Nature 461, 361–366. 10.1038/nature08319.

31. McDowell, M.A., Heimes, M., Fiorentino, F., Mehmood, S., Farkas, A., Coy-Vergara, J., Wu, D., Bolla, J.R., Schmid, V., Heinze, R., et al. (2020). Structural Basis of Tail-Anchored Membrane Protein Biogenesis by the GET Insertase Complex. Mol Cell 80, 72–86 e77. 10.1016/j.molcel.2020.08.012.

32. Coy-Vergara, J., Rivera-Monroy, J., Urlaub, H., Lenz, C., and Schwappach, B. (2019). A trap mutant reveals the physiological client spectrum of TRC40. J Cell Sci 132. 10.1242/jcs.230094.

33. Bozkurt, G., Stjepanovic, G., Vilardi, F., Amlacher, S., Wild, K., Bange, G., Favaloro, V., Rippe, K., Hurt, E., Dobberstein, B., and Sinning, I. (2009). Structural insights into tail-anchored protein binding and membrane insertion by Get3. Proc Natl Acad Sci U S A 106, 21131–21136. 10.1073/pnas.0910223106.

34. Hu, J., Li, J., Qian, X., Denic, V., and Sha, B. (2009). The crystal structures of yeast Get3 suggest a mechanism for tail-anchored protein membrane insertion. PLoS One 4, e8061. 10.1371/journal.pone.0008061.

35. Stefer, S., Reitz, S., Wang, F., Wild, K., Pang, Y.Y., Schwarz, D., Bomke, J., Hein, C., Lohr, F., Bernhard, F., et al. (2011). Structural basis for tail-anchored membrane protein biogenesis by the Get3-receptor complex. Science 333, 758–762. 10.1126/science.1207125.

36. Yamagata, A., Mimura, H., Sato, Y., Yamashita, M., Yoshikawa, A., and Fukai, S. (2010). Structural insight into the membrane insertion of tail-anchored proteins by Get3. Genes Cells 15, 29–41. 10.1111/j.1365-2443.2009.01362.x.

37. Mariappan, M., Li, X., Stefanovic, S., Sharma, A., Mateja, A., Keenan, R.J., and Hegde, R.S. (2010). A ribosome-associating factor chaperones tail-anchored membrane proteins. Nature 466, 1120–1124. 10.1038/nature09296.

38. Sontag, E.M., Samant, R.S., and Frydman, J. (2017). Mechanisms and Functions of Spatial Protein Quality Control. Annu Rev Biochem 86, 97–122. 10.1146/annurev-biochem-060815-014616.

39. Pfaff, J., Rivera Monroy, J., Jamieson, C., Rajanala, K., Vilardi, F., Schwappach, B., and Kehlenbach, R.H. (2016). Emery-Dreifuss muscular dystrophy mutations impair TRC40-mediated targeting of emerin to the inner nuclear membrane. J Cell Sci 129, 502–516. 10.1242/jcs.179333.

40. Sheth, U., and Parker, R. (2003). Decapping and decay of messenger RNA occur in cytoplasmic processing bodies. Science 300, 805–808. 10.1126/science.1082320.

41. Aulas, A., Caron, G., Gkogkas, C.G., Mohamed, N.V., Destroismaisons, L., Sonenberg, N., Leclerc, N., Parker, J.A., and Vande Velde, C. (2015). G3BP1 promotes stress-induced RNA granule interactions to preserve polyadenylated mRNA. J Cell Biol 209, 73–84. 10.1083/jcb.201408092.

42. Kageyama, S., Gudmundsson, S.R., Sou, Y.S., Ichimura, Y., Tamura, N., Kazuno, S., Ueno, T., Miura, Y., Noshiro, D., Abe, M., et al. (2021). p62/SQSTM1-droplet serves as a platform for autophagosome formation and anti-oxidative stress response. Nat Commun 12, 16. 10.1038/s41467-020-20185-1.

43. Imamura, H., Nhat, K.P., Togawa, H., Saito, K., Iino, R., Kato-Yamada, Y., Nagai, T., and Noji, H. (2009). Visualization of ATP levels inside single living cells with fluorescence resonance energy transfer-based genetically encoded indicators. Proc Natl Acad Sci U S A 106, 15651–15656. 10.1073/pnas.0904764106.

44. Johnston, J.A., Ward, C.L., and Kopito, R.R. (1998). Aggresomes: a cellular response to misfolded proteins. J Cell Biol 143, 1883–1898. 10.1083/jcb.143.7.1883.

45. Bressan, C., Pecora, A., Gagnon, D., Snapyan, M., Labrecque, S., De Koninck, P., Parent, M., and Saghatelyan, A. (2020). The dynamic interplay between ATP/ADP levels and autophagy sustain neuronal migration in vivo. Elife 9. 10.7554/eLife.56006.

46. Zhang, J., Goliwas, K.F., Wang, W., Taufalele, P.V., Bordeleau, F., and Reinhart-King, C.A. (2019). Energetic regulation of coordinated leader-follower dynamics during collective invasion of breast cancer cells. Proc Natl Acad Sci U S A 116, 7867–7872. 10.1073/pnas.1809964116.

47. Jang, S., Nelson, J.C., Bend, E.G., Rodriguez-Laureano, L., Tueros, F.G., Cartagenova, L., Underwood, K., Jorgensen, E.M., and Colon-Ramos, D.A. (2016). Glycolytic Enzymes Localize to Synapses under Energy Stress to Support Synaptic Function. Neuron 90, 278–291. 10.1016/j.neuron.2016.03.011.

48. Houck, S.A., Singh, S., and Cyr, D.M. (2012). Cellular responses to misfolded proteins and protein aggregates. Methods Mol Biol 832, 455–461. 10.1007/978-1-61779-474-2_32.

49. Jeng, W., Lee, S., Sung, N., Lee, J., and Tsai, F.T. (2015). Molecular chaperones: guardians of the proteome in normal and disease states. F1000Res 4. 10.12688/f1000research.7214.1.

50. Escusa-Toret, S., Vonk, W.I., and Frydman, J. (2013). Spatial sequestration of misfolded proteins by a dynamic chaperone pathway enhances cellular fitness during stress. Nat Cell Biol 15, 1231–1243. 10.1038/ncb2838.

51. Kaganovich, D., Kopito, R., and Frydman, J. (2008). Misfolded proteins partition between two distinct quality control compartments. Nature 454, 1088–1095. 10.1038/nature07195.

52. Chernoff, Y.O., Lindquist, S.L., Ono, B., Inge-Vechtomov, S.G., and Liebman, S.W. (1995). Role of the chaperone protein Hsp104 in propagation of the yeast prion-like factor [psi+]. Science 268, 880–884. 10.1126/science.7754373.

53. Doyle, S.M., Genest, O., and Wickner, S. (2013). Protein rescue from aggregates by powerful molecular chaperone machines. Nat Rev Mol Cell Biol 14, 617–629. 10.1038/nrm3660.

54. Mogk, A., Bukau, B., and Kampinga, H.H. (2018). Cellular Handling of Protein Aggregates by Disaggregation Machines. Mol Cell 69, 214–226. 10.1016/j.molcel.2018.01.004.

55. Tittelmeier, J., Sandhof, C.A., Ries, H.M., Druffel-Augustin, S., Mogk, A., Bukau, B., and Nussbaum-Krammer, C. (2020). The HSP110/HSP70 disaggregation system generates spreading-competent toxic alpha-synuclein species. EMBO J 39, e103954. 10.15252/embj.2019103954.

56. Ulrich, K., Schwappach, B., and Jakob, U. (2020). Thiol-based switching mechanisms of stress-sensing chaperones. Biol Chem. 10.1515/hsz-2020-0262.

57. Jain, S., Wheeler, J.R., Walters, R.W., Agrawal, A., Barsic, A., and Parker, R. (2016). ATPase-Modulated Stress Granules Contain a Diverse Proteome and Substructure. Cell 164, 487–498. 10.1016/j.cell.2015.12.038.

58. Hyttinen, J.M., Amadio, M., Viiri, J., Pascale, A., Salminen, A., and Kaarniranta, K. (2014). Clearance of misfolded and aggregated proteins by aggrephagy and implications for aggregation diseases. Ageing Res Rev 18, 16–28. 10.1016/j.arr.2014.07.002.

59. Rahman, M.A., Rahman, M.D.H., Mamun-Or-Rashid, A.N.M., Hwang, H., Chung, S., Kim, B., and Rhim, H. (2022). Autophagy Modulation in Aggresome Formation: Emerging Implications and Treatments of Alzheimer’s Disease. Biomedicines 10. 10.3390/biomedicines10051027.

60. Buchner, J., Grallert, H., and Jakob, U. (1998). Analysis of chaperone function using citrate synthase as nonnative substrate protein. Methods Enzymol 290, 323–338. 10.1016/s0076-6879(98)90029-5.

61. Jakob, U., Gaestel, M., Engel, K., and Buchner, J. (1993). Small heat shock proteins are molecular chaperones. J Biol Chem 268, 1517–1520.

62. Kiianitsa, K., Solinger, J.A., and Heyer, W.D. (2003). NADH-coupled microplate photometric assay for kinetic studies of ATP-hydrolyzing enzymes with low and high specific activities. Anal Biochem 321, 266–271. 10.1016/s0003-2697(03)00461-5.

63. Leichert, L.I., Gehrke, F., Gudiseva, H.V., Blackwell, T., Ilbert, M., Walker, A.K., Strahler, J.R., Andrews, P.C., and Jakob, U. (2008). Quantifying changes in the thiol redox proteome upon oxidative stress in vivo. Proc Natl Acad Sci U S A 105, 8197–8202. 10.1073/pnas.0707723105.

